# Critical *cis*-parameters influence STructure Assisted RNA Translation (START) initiation on non-AUG codons in eukaryotes

**DOI:** 10.1101/2024.02.23.581729

**Authors:** Antonin Tidu, Fatima Alghoul, Laurence Despons, Gilbert Eriani, Franck Martin

## Abstract

In eukaryotes, translation initiation is a highly regulated process, which combines *cis-* regulatory sequences located on the messenger RNA along with *trans-*acting factors like eukaryotic initiation factors (eIF). One critical step of translation initiation is the start codon recognition by the scanning 43S particle, which leads to ribosome assembly and protein synthesis. In this study, we investigated the involvement of secondary structures downstream the initiation codon in the so-called START (Structure-Assisted RNA translation) mechanism on AUG and non-AUG translation initiation. The results demonstrate that downstream secondary structures can efficiently promote non-AUG translation initiation provided that they are stable enough to stall a scanning 43S particle and that they are located at an optimal distance from this non-AUG codon to trigger and stabilize the codon-anticodon base-pairing in the P site. The required stability of the downstream structure for efficient translation initiation varies in distinct cell types. We extended this study to genome-wide analysis of the *Homo sapiens* alternative translation initiation sites and discovered 556 of them starting with an AUG and 506 starting with a non-AUG that contained a downstream RNA structure at an optimal distance and with a predicted stability of at least -15 kcal/mol. We validated the impact of these structures on translation initiation for several selected uORFs.

## INTRODUCTION

In eukaryotes, translation initiation is a highly regulated process, which combines *cis-* regulatory sequences located on the messenger RNA along with *trans-*acting factors like eukaryotic initiation factors (eIF) ^1^. The canonical translation initiation mechanism starts with the recognition of the 5’ m^7^G cap by eIF4E, which is associated with eIF4G and eIF4A to form the eIF4F complex. The interaction between eIF4G and 43S-bound eIF3 enables the recruitment of the 43S particle on the m^7^G cap. Then, the 43S particle scans the 5’UTR thanks to the ATP-dependent RNA helicase activity of eIF4A, which is enhanced through its interactions with eIF4G, eIF4B (and/or eIF4H) and eIF3 ^2,3^. During scanning, the 43S particle is maintained in a so-called open conformation by the combined actions of eIF1A and eIF1 which destabilize the codon-anticodon interaction in the P-site, inhibit spontaneous hydrolysis of eIF2-bound GTP ^4^ and therefore prevent ribosomal subunits joining ^5–7^. The codon-anticodon interaction involves the nucleotide triplet from the codon being analyzed in the P-site and the anticodon of the initiator Met-tRNA^Met^_i_ which is bound to eIF2-GTP in the 43S particle ^8^. When a stable enough codon-anticodon base-pairing is established during scanning, it displaces eIF1 from the P-site and triggers the switch from the open to the close conformation of the 43S. This results in the activation of the GTPase activity of eIF2 by eIF5 and in the subsequent hydrolysis of eIF2-bound GTP into GDP ^9^. As GTP hydrolysis goes on, eIF2 loses its affinity for the Met-tRNA^Met^_i_ and is released from the 48S particle ^10^. Subunits joining is mediated by eIF5B ^11^ and its interaction with eIF1A ^12^, which contributes to the release eIF2, eIF1, eIF1A and eIF5B from the initiation complex through the hydrolysis of eIF5B-bound GTP. The assembled 80S ribosome then starts protein synthesis by incorporating the first amino-acyl-tRNA in its vacant A-site.

Start-codon selection by the scanning 43S particle is the most critical step of initiation as it determines the amino-acid sequence of the protein that will be synthesized, and therefore has drastic consequences for its function(s). Incorrect start site selection can result in N-terminal extensions or deletions, leading to erroneous protein isoforms, or to out-of-frame protein synthesis, leading to an unrelated and non-functional translation product. Among several parameters, the most critical feature of start codon selection is the stability of the codon-anticodon interaction. Start-codon selection stringency is mediated by eIF1 and eIF1A which prevent unstable codon-anticodon interactions like those involving non-cognate codons ^5–7^. However, even AUG codons that establish a perfect Watson-Crick base-pairing with the anticodon loop of the Met-tRNA^Met^_i_ are sometimes not recognized as the initiation codon. Moreover, several studies ^13,14^ along with ribosome profiling data ^15–17^ suggest that non-AUG codons can be used for translation initiation in eukaryotes. Therefore start-codon selection is influenced by additional *trans-* and *cis-*acting factors ^1^. Several reports, starting with the seminal studies by Marylin Kozak ^18,19^, suggest that the start codon nucleotide context plays a major role on start codon recognition efficiency. Distal secondary structures elements have also been shown to modulate start codon selection stringency ^20^. In a previous study, we showed that translation of C9ORF72 transcripts into toxic poly-dipeptides is initiated on a CUG codon that is upstream of very stable RNA structures that contain G-quartets ^14^. More broadly, it has been shown that the impairment of the RNA helicase Ded1p is linked to increased uORFs translation in yeast ^21^. Both observations suggest the involvement of downstream RNA secondary structures in start-codon selection. Similarly, proteins bound to an RNA element downstream of an initiation site can promote start codon recognition. An example is the Drosophila protein Sex-Lethal (SXL) that promotes uORF translation of male specific lethal 2 mRNA by stalling 43S complexes upstream the main open reading frame ^22,23^. Based on these observations, we previously described a mechanism of translation initiation called STructure Assisted RNA Translation or START ^24,25^. START initiation relies on the presence of a downstream structure that enhances translation initiation on both AUG and non-AUG codons by stalling the scanning 43S particle with those codons in the P-site, leading to 80S assembly and protein synthesis on these sub-optimal initiation codons.

In this work, we aimed to determine to what extent non-AUG translation initiation relies on a START mechanism. We have precisely investigated three critical parameters for START in the presence of a downstream RNA structure to the initiation codon: the distance between the structure and the P-site, the structure stability, and the type of the start codon. We show that downstream secondary structures can efficiently promote non-AUG translation initiation if they are stable enough to stall a scanning 43S particle and located at an optimal distance from this non-AUG codon to stabilize the codon-anticodon base pairing in the P site. In addition, the efficiency of START relies on the start codon type along with its nucleotide environment and is also highly influenced by *trans*-acting factors. We have previously shown the broad presence of downstream RNA structures in annotated main open reading frames of various organisms ^25^. Here, we further extended this study by analysing genome-wide alternative translation initiation sites in human transcripts that were previously determined by ribosome profiling ^17^ and found the broad presence of moderately stable downstream RNA structures within the distance range we have determined being optimal for START. Functional analysis of some examples confirmed the role of downstream structures on translation efficiency at these alternative translation initiation sites.

## MATERIAL AND METHODS

### Oligonucleotides information

All the oligonucleotides’ sequences used in this study are provided in supplementary material (Supplementary Table 1). Oligonucleotides were purchased from Integrated DNA Technologies company. In the text, oligonucleotides are referred to by their red numbers in the Supplementary Table 1.

### Rabbit reticulocyte lysate

Untreated rabbit reticulocyte lysate (RRL) was prepared as previously described ^26^.

### HEK293FT and SH-SY5Y cell lysates from cells cultured in physiological conditions

HEK293FT cells were seeded at 0.03 x10^6^ cells/cm² and grown for 2 days at 37°C in a 5% CO_2_ humidified atmosphere. The culture medium is Gibco Dulbecco’s Modified Eagle Medium + GlutaMAX (Life Technologies) which is supplemented with 10% of inactivated fetal bovine serum (Life Technologies). Neuroblastoma SH-SY5Y cells were cultured in the same conditions and the same medium but supplemented with 1 mM of non-essential amino acids (Gibco). The total culture surface was 4500 cm², leading approx. to 200 to 400 x 10^6^ cells before harvesting. Cells were harvested at room temperature in their culture medium by centrifugation (300g) at 4°C and washed two times with a cold buffer containing 20 mM HEPES–KOH pH 7.5, 100 mM potassium acetate, 2 mM magnesium acetate, 1 mM DTT. After washing, the cell pellets were resuspended in the same buffer supplemented or not (if TEV protease is used, see later) with 1X Halt™ Protease Inhibitor Cocktail EDTA-free (Thermo Scientific™) to reach a concentration of 50-100 x10^6^ cells/mL. Cells were lysed by nitrogen cavitation with a Cell Disruption Bomb (Parr Instrument Company) after a one-hour incubation under a pressure of 30 bars at 4°C. The lysate was cleared by centrifugations at 10,000g at 4°C, aliquoted, flash-frozen in liquid nitrogen and stored at −80°C. Total protein concentration was determined by Bradford assay (Biorad).

### HEK293FT cell lysates from cells cultured in stress conditions

To prepare cell-free translation extracts from HEK293FT cells cultured in stress conditions, the same procedure was used except that the culture medium was supplemented with 5 mM DTT 3 hours before harvesting to induce endoplasmic reticulum (ER) stress, which triggers the unfolded protein response (UPR) ^27,28^. This pathway leads in particular to the phosphorylation of the eIF2 α-subunit by the protein kinase R-like endoplasmic reticulum kinase (PERK) ^29,30^.

### Measurements of eIF2 α-subunit phosphorylation by Western Blot

Aliquots of *in vitro* translation reactions were run on 12% SDS-PAGE. Proteins were transferred on PVDF membranes (Immobilon®-P Transfer Membrane) for 1h at 10V. Membranes were saturated with [PBS 1X, Tween20 0.5%, BSA 50mg/mL]. The primary antibodies (@eIF2α: #9722 and @eIF2α-phosphorylated: #3597, Cell Signaling Technology) were diluted ten thousand times in PBS 1X, Tween 20 0.5%, BSA 50 mg/mL and hybridized overnight at 4°C. Membranes were then washed three times in PBS 1X, Tween20 0.5% before the 2-hour hybridization of the secondary antibody (@rabbit A120-101-P, Bethyl Laboratories) which was diluted ten thousand times in PBS 1X, Tween20 0.1%. Membranes were finally washed three times in PBS 1X, Tween20 0.1% before the addition of the ECL substrate (Clarity Western ECL Substrate #1705061). Chemiluminescence signals were measured with a Biorad ChemiDoc (MP) apparatus and resulting TIFF images were analyzed with ImageJ software.

### Reporter RNA synthesis for *in vitro* translation

All plasmids containing the 5’UTR and protein reporter sequences of interest were prepared using the NEBuilder^®^ HiFi DNA Assembly Cloning Kit (#E5520S). Site-directed mutagenesis was performed with NEB Q5® Site Directed Mutagenesis Kit (#E0554S). Then, the corresponding T7-5’UTR-reporter DNA construct was PCR-amplified using forward primer n°1 (capped reporters) or n°2 (IGR reporters) and reverse primer n°3 (Supplementary Table 1). Corresponding RNA reporters (Supplementary File 1) were synthesized in a 100 µL *in vitro* transcription reaction using 0.1 µM of T7-DNA template, 5 mM Tris-HCl pH 8, 30 mM MgCl_2_, 1 mM spermidine, 5 mM DTT, 0.01% Triton X-100, 5 mM of each ribonucleotide (ATP, CTP, GTP, UTP pH 7.5), 0.5 U/µL RNase inhibitor (Promega) and 0.125 mg/mL of home-made recombinant T7 RNA polymerase. The reaction was incubated for 1h at 37°C. For co-transcriptional capping, the reaction was started in the absence of GTP and in the presence of 0.5 mM of anti-reverse m^7^G-cap analog (NEB #S1411) for 10 min. GTP was then gradually added up to 5 mM at 1h incubation. After 1h incubation, 0.02 mg/mL pyrophosphatase (Merck) was added and after 30 min, the DNA template was degraded by 1h incubation with 0.2U/µL DNaseI (Roche). Transcription products were loaded on a 1 mL G-25 Superfine Sephadex column (Cytiva) and the resulting eluate was phenol-extracted and ethanol-precipitated. The resulting RNA pellets were resuspended in 30 µL milli-Q water, frozen, and quantified by 260 nm absorbance measurement. All reporter RNA sequences used in this work are provided in Supplementary File 1.

### *In vitro* translation assays

#### RRL

the final composition in the reaction was 50% RRL, 100 mM potassium acetate, 1 mM magnesium acetate, 0.1 mM of all amino acids but methionine, 0.5 U/µL RNase inhibitor (Promega), 0.125 µCi/µL ^35^S-methionine (Perkin-Elmer) and 0.2 µM reporter RNA in a total volume of 20 µL. The reaction was incubated for 1h at 30°C or 2h at 30°C when TEV protease was used for co-translational cleavage.

#### HEK293FT and SH-SY5Y extracts

the final composition of the translation mixture was 15% cell extract (1.8 µg/µL total protein), 100 mM potassium acetate, 1 mM magnesium acetate, 0.1 mM of all amino acids but methionine, 20 mM HEPES-KOH pH 7.5, 0.5 mM spermidine, 1 mM DTT, 0.8 mM ATP, 0.1 mM GTP, 8 mM phospho-creatine, 0.1 µg/µL creatine phospho-kinase, 0.5 U/µL RNase inhibitor (Promega), 0.125 µCi/µL ^35^S-methionine (Perkin-Elmer) and 0.2 µM reporter RNA in a total volume of 20 µL. The reaction was incubated for 1h at 30°C or 2h at 30°C when TEV protease was used for co-translational cleavage.

#### Co-translational cleavage of *in vitro* synthesized reporter proteins with TEV protease

cleavage was triggered by the addition of recombinant home-made TEV protease to *in vitro* translation reactions at a final concentration of 0.035 µg/µL.

### Assessment of *in vitro* translation products levels

#### ^35^S-signal acquisition

5 µL of *in vitro* translation reactions were analyzed by 12% SDS-PAGE. ^35^S-labelled translation products bands were quantified with a phosphorimager (Typhoon FLA 7000). Image analysis was conducted with ImageJ software using the resulting TIFF files.

#### Luciferase assay

the amount of *in vitro* synthetized Renilla luciferase was assessed upon injection of 100 µL coelenterazine 0.25 µmol/mL (Synchem) into 10 µL of *in vitro* translation reaction using a luminometer (Varioskan LUX, Thermo Scientific™). Subsequent photon emission was measured for 10 seconds with the default measurement parameters.

Raw luminescence and ^35^S-signal were processed using a self-made Python script. For luminescence signals, as each sample was assessed in triplicates, descriptive (mean, standard deviation) and inferential statistics (Student p-values) of the ratio of sample A (triplicates a1, a2, a3) over sample B (triplicates b1, b2, b3) were determined using the nine possible ratios between (a1, a2, a3) and (b1, b2, b3). For ^35^S-signal, as signals were normalized within each 3 gels independently, this resulted in 3 possible ratios for assessing the statistics of A/B. Statistical tests and annotations were performed with the statannotations package (v 0.4.4.).

### Predictions and free energy calculations of RNA secondary structures

These calculations were conducted using the Mfold V2.3 for RNA ^31^ on the various a11 structure mutants used in this work. Constraints were implemented according to the probing data that enabled us to draw the 2D model of the wild-type structure ^32^.

### Automated secondary structures predictions in transcripts

RNA transcript sequence recovery from human smORF data ^17^ was performed using two homemade Python scripts available as supplementary data. They must be executed in the following order: (1) format_smORFs_seq.py, (2) select_smORF_sequences.py. Nucleotide sequences between the +16 and +65 positions relative to the initiator codon of smORFs were extracted. When the smORF coding sequences were shorter than 65 bases, BLASTN searches ^33^ were run against local human RNA databases to find the corresponding transcripts. Two additional scripts were used to search and then analyze smORF structures: (3) search_2Dstruct_in_smORFs.py, (4) analyse_2Dstruct_in_smORFs.py. Scripts 3 and 4 are based on scripts used in a previous study ^25^ and modified to suit the prediction or analysis of secondary structures in the +16 to +65 region of the smORF transcripts. The structure prediction algorithm implemented in these scripts is RNAfold from the ViennaRNA package 2 ^34^.RNAfold program calculates the minimum free energy (MFE) of RNA secondary structures such as stem-loops and G-Quadruplexes. The Python module named toolbox.py is imported by some of these scripts (Supplementary File 4: Python_scripts.zip).

### Gene Ontology (GO) terms analysis

GO-terms analysis was performed using the relevant functions from the GOATools package, version 1.3.11 ^35^. For GO-terms enrichment analysis, p-values from Fisher exact tests were corrected using the Holm method. The gene ontology file (go-basic.obo) was downloaded from <u>Download ontology</u> (geneontology.org) in its latest version to date (release from 15/11/2023). Human annotation file (goa_human.gaf) was downloaded from <u>Download Annotations | Gene Ontology Consortium</u> in its latest version to date (Panther version v.17.0, GO version 2023-10-09).

## RESULTS

### Experimental strategy to investigate the impact of secondary structures in the coding sequence

To test the impact of several critical parameters for START, we synthesized a set of reporter mRNAs that encode the Renilla luciferase fused to variable N-terminal peptides resulting from the translation of various RNA structures downstream of the start codon (Figure 1). To avoid any putative artifacts due to variable N-terminal sequences that would influence the luciferase activity of the Renilla protein, we introduced a TEV protease cleavage upstream of the Renilla coding sequence. The quantification of luciferase activity was performed after TEV cleavages which yield identical Renilla proteins produced by all the reporter mRNAs. The sequences of all the reporter RNA used in this study are shown in Supplementary File 1. More precisely, these reporter mRNAs feature a linear CAA-rich 5’UTR, to minimize the effect of structures on 5’-3’ scanning, followed by a start codon of interest. These reporter mRNAs were co-transcriptionally m^7^G-capped and their translation efficiency was determined by Renilla luciferase activity measurements. With these reporters, we investigated the influence of both the location and the structure stability on translation initiation efficiency, as well as the nature of the start codon and its nucleotide context. We confirmed that the cell-free translation extracts used in this study are cap-dependent (Supplementary Figure 1).

**Figure 1:**
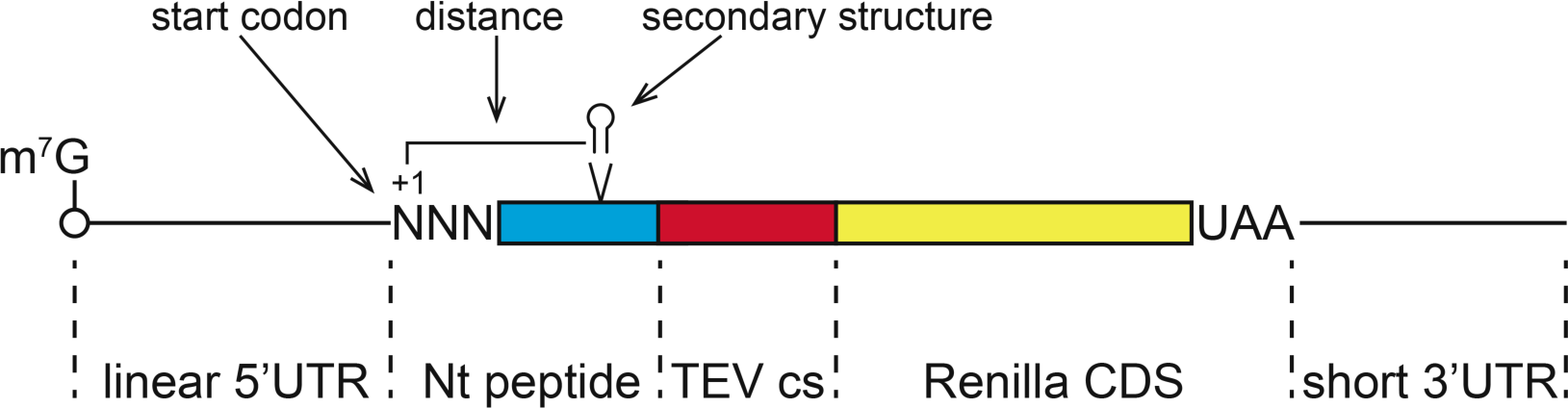
Schematic representation of the RNA reporters used in the study. The three *cis-*acting parameters studied are (i) the distance between a downstream structure and the start codon, (ii) the stability of the downstream structure, (iii) the nature of the start codon. TEV cs: Tobacco Etch Virus protease cleavage site, Nt peptide: N-terminal fusion peptide, Renilla CDS: Renilla luciferase coding sequence.

As a model secondary structure, we used a 50-nucleotide GC-rich secondary structure from the 5’UTR of Hox a11 mRNA. We have previously determined its secondary structure by probing and shown that this structure is very stable (ΔG = -21,6 kcal/mol) as it consists of 16 G-C base pairs. Importantly, the a11 structure can stall a scanning complex ^32^.

### The optimal position of the secondary structure for translation on a CUG codon ranges from +23 to +26

To determine the optimal distance between the a11 secondary structure and the start codon, *in vitro* translation assays using untreated rabbit reticulocyte lysate were conducted with reporter mRNAs containing an AUG or a CUG start codon with the a11 secondary structure located at positions ranging from +11 to +35 (Figure 2A). AUG translation was relatively insensitive to the presence of the secondary structure, whatever its position (Figure 2B-C). On the other hand, CUG translation varied significantly depending on the position of the secondary structure (Figure 2C). It is of note significantly impaired when the structure is located at positions +11 or +35. In addition, a stronger increase in relative translation efficiency is observed for CUG compared to AUG at position +26, indicating the optimal position of the structure would be close to +26 for CUG translation initiation. We then determined the optimal position of the structure for CUG translation (Figure 3A). Translation efficiency decreases when the structure is located before +17 or beyond +32 from the CUG codon (Figure 3B-C). Overall, these results show that while AUG initiation is rather insensitive to downstream secondary structures located between +11 and +35, CUG initiation requires a structure located between +23 and +26.

**Figure 2:**
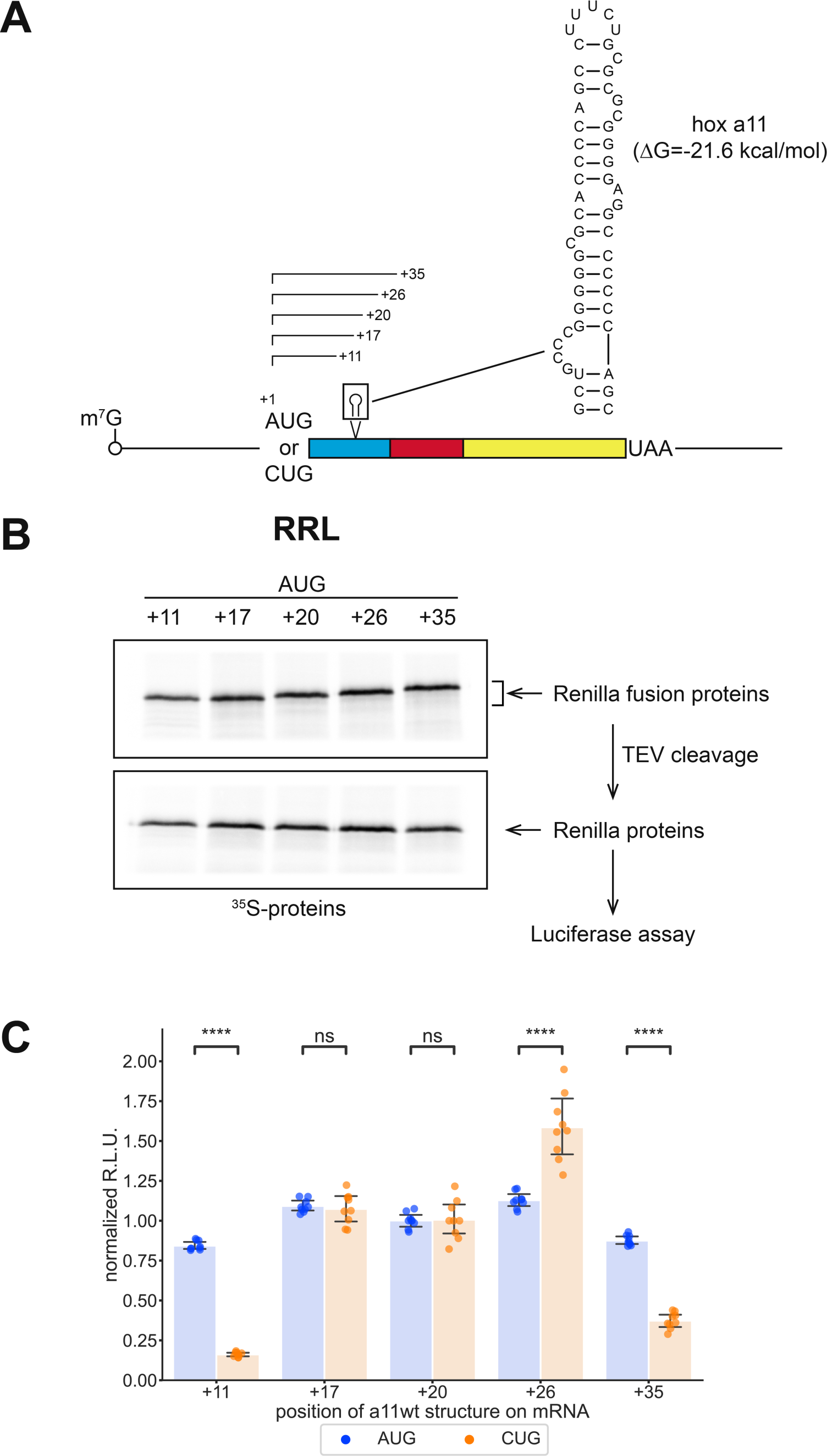
Translation initiation on CUG is sensitive to the position of the downstream secondary structure. **A.** Schematic representation of the RNA reporters highlighting the two AUG or CUG initiation codons used and the wild-type a11 hairpin, which is inserted at +11, +17, +20, +26 or +35 positions. **B.** Representative SDS-PAGE of *in vitro* translation products obtained with AUG reporters in RRL. The upper panel shows the ^35^S-labelled products before TEV cleavage, the lower panel shows the ^35^S-labelled products after TEV cleavage. Proteins produced by TEV cleavage were used for Luciferase assays. **C.** Quantification of Renilla luciferase luminescence produced in RRL from a11-RNA reporters initiating with an AUG codon (blue) or with a CUG codon (orange). Relative luminescence units (R.L.U.) were normalized to those obtained with the +20 AUG reporter or CUG reporter. The 9 possible normalized ratios (a/b) values that can be obtained from 3 independent replicates are shown as blue (with AUG) and orange (with CUG) dots, while bar heights show mean normalized ratios. Error bars represent the 99% confidence interval calculated from the 9 normalized ratios using the t-distribution. p-values were calculated using a student t-test for independent samples. *: 0.01 < p < 0.05, **: 0.001 < p < 0.01, ***: 0.0001 < p < 0.001, ****: p < 0.0001

**Figure 3:**
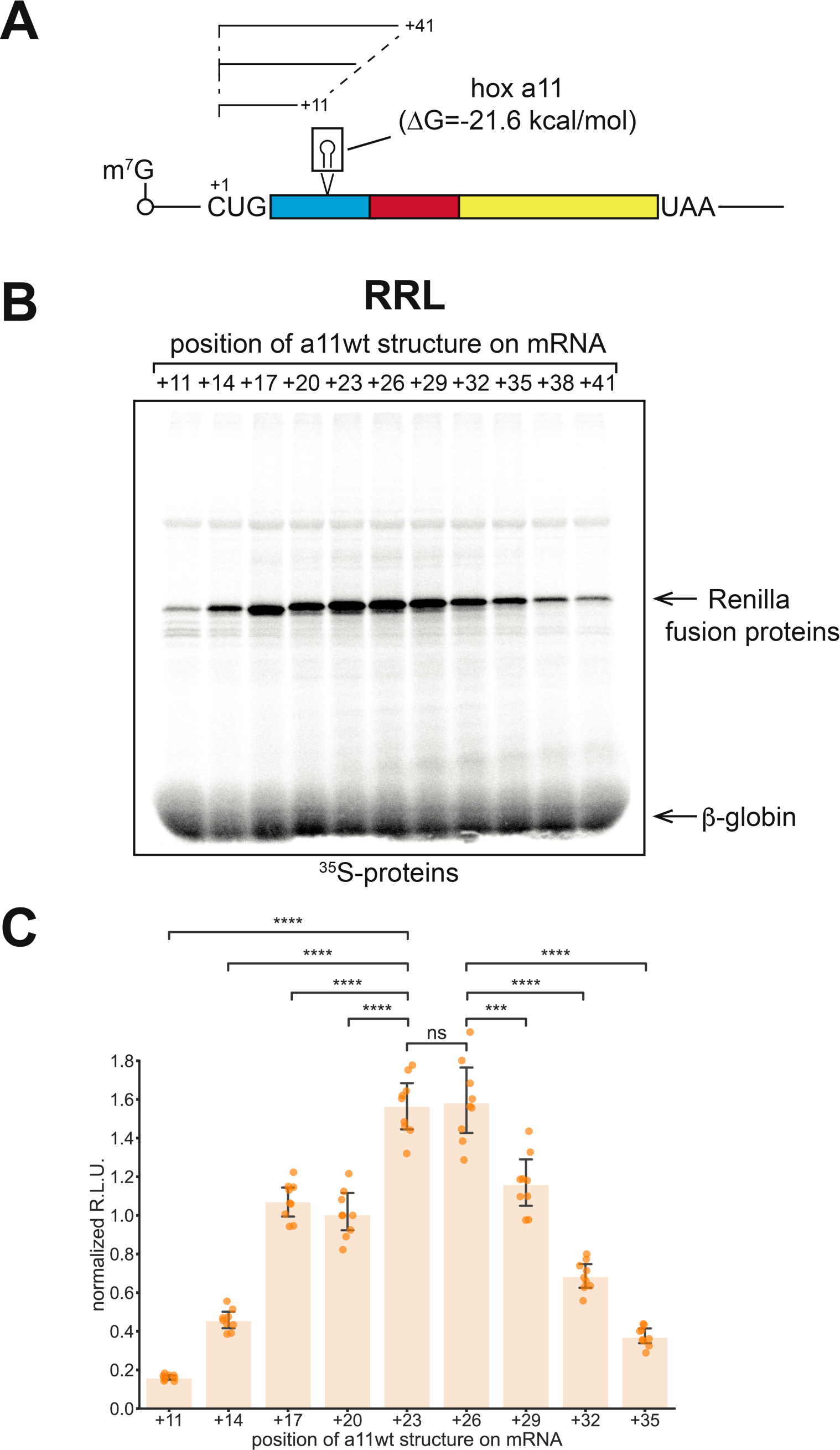
The optimal position of the secondary structure for initiation on a CUG codon is between +23 to +26. **A.** Schematic representation of the RNA reporters initiating on a CUG codon upstream of the wild-type a11 hairpin inserted at positions +11, +17, +20, +23, +26, +29, +32, +35, +38 or +41. **B.** Representative SDS-PAGE of *in vitro* ^35^S radiolabelled translation products obtained with AUG reporters in RRL. **C.** Quantification of Renilla luciferase luminescence produced in RRL from a11 RNA reporters initiating with a CUG codon. Relative luminescence units (R.L.U.) were normalized to those obtained with the +20 CUG reporter. Bar heights show the mean values with standard deviations. P-values were calculated using a student t-test for independent samples. *: 0.01 < p < 0.05, **: 0.001 < p < 0.01, ***: 0.0001 < p < 0.001, ****: p < 0.0001.

### The optimal stability of the secondary structure is influenced by the cell type

Next, we examined the required stability of the secondary structure for efficient translation initiation on a CUG codon. We used reporters with an AUG as a control and a CUG start codon upstream of the a11 secondary structure that is localized at the rather optimal position +20, as determined in the previous section. The inserted a11 structures contain mutations that progressively decrease the number of G-C base pairs, resulting in predicted stabilities ranging from -21.6 to -4 kcal/mol (Figure 4A). Silent mutations were used to avoid any putative co-translational side effects due to amino acid substitutions, except for the -4 kcal/mol mutant structure that contains mutations leading to 3 amino acids changes in the N-terminal fusion of the translated reporter protein. However, since TEV cleavage was performed co-translationally, all the translated mRNAs produced the same Renilla luciferase protein. We therefore eliminated potential N-terminal induced co-translational artifacts on the activity of the Renilla luciferase protein.

**Figure 4:**
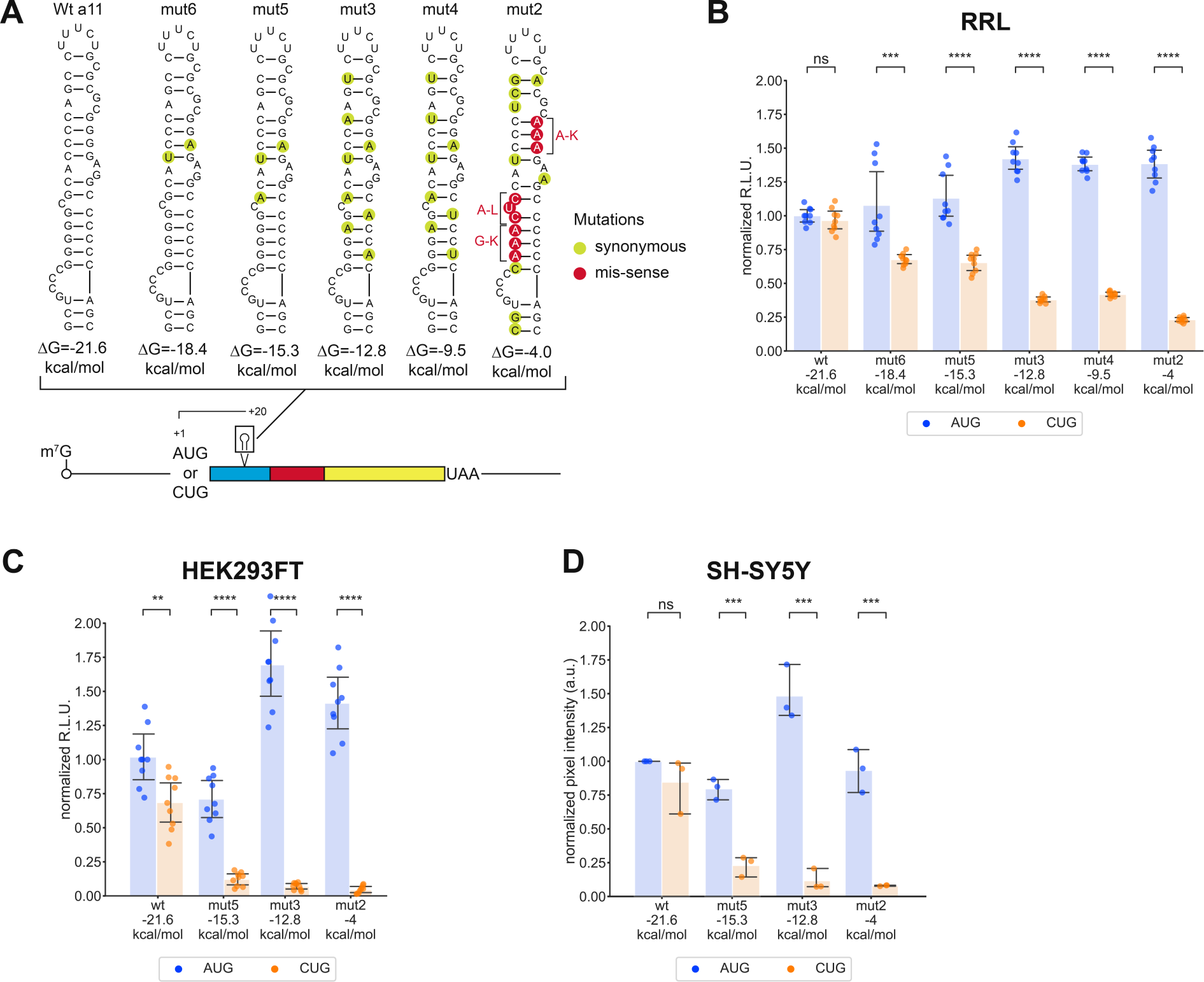
Secondary structure stability determines translation initiation efficiency in different cell extracts. **A.** Schematic representation of the RNA reporter and sequences of the stem-loops used during the study. The initiation codons were AUG or CUG. Wild-type and mutant a11 hairpins were inserted at +20 position. Silent mutations introduced to lower the a11 structure stability are shown in green circles whereas missense mutations are shown in red circles. **B.** Quantification of Renilla luciferase luminescence produced in RRL from variable a11 RNA reporters initiating with AUG or CUG codon. Relative luminescence units (R.L.U.) were normalized to those obtained with the reporter containing the wild-type a11 structure and AUG codon. Bar heights show the mean values with standard deviations indicated. P-values were calculated using a student t-test for independent samples. *: 0.01 < p < 0.05, **: 0.001 < p < 0.01, ***: 0.0001 < p < 0.001, ****: p < 0.0001. **C.** Quantification of Renilla luciferase luminescence produced in HEK293FT cell extracts from variable a11 RNA reporters initiating with AUG or CUG codon. Results are presented as in B. **D.** Quantification of ^35^S incorporation in Renilla luciferase proteins produced in SH-SY5Y cell extracts. Results are presented as in B and C.

We first confirmed with rabbit reticulocyte lysates that AUG-reporters translation does not rely on downstream structures (Figure 4B). On the contrary, initiation on the CUG codon decreases as soon as secondary structure stability is below -21 kcal/mol. This shows the importance of a downstream secondary structure with a stability of at least -15 kcal/mol for the initiation of translation on a CUG codon using these experimental conditions.

We have developed and characterized cell-free *in vitro* translation extracts from human embryonic kidney (HEK293FT) (Supplementary Figures 1 and 2) and neuroblastoma (SH-SY5Y) cell lines, which makes it possible to analyze START in the presence of cell-type-specific *trans-*acting factors. Translation of the reporter RNAs using these human cell extracts resulted in the same trend as with RRL (Figure 4C-D). However, the minimal stability required to reach efficient translation initiation on a CUG codon is different with these extracts. Indeed, initiation on a CUG codon strictly requires an a11 secondary structure with a stability of -21.6 kcal/mol. When the a11 structure stability is diminished, translation from the CUG is drastically reduced, in contrast to the gradual reduction observed with RRL. This indicates that HEK293FT and SH-SY5Y extracts are much more sensitive to the structure stability for START than RRL. This difference with RRL might be attributed to a distinct content of RNA helicases or RNA binding proteins that could unwind or stabilize the downstream structure required for CUG initiation.

### The START mechanism alone does not promote translation initiation on all AUG-like codons

Next, we extended these experiments to other AUG-like codons (CUG, GUG, UUG, ACG and AUC) with the -15 kcal/mol a11 structure located at position +20 in RRL or the wt a11 structure (-21.6 kcal/mol) for experiments with HEK293FT extracts (Figure 5A). Overall, the AUG-like codons are generally less efficiently recognized by the ribosome. In addition, all AUG-like codons are not equivalently recognized and follow the hierarchy ACG>UUG>AUC>GUG≈0. The result obtained with GUG must however be nuanced, as the inefficiency of GUG initiation is most likely linked to the reporter RNA sequence around the initiation site. When a GUG is introduced in the initiation context of these reporters, it introduces an upstream out-of-frame *AUG* that overlaps the <u>GUG</u> (CGUA*AU<u>G</u>*<u>UG</u>GAC). Such a nucleotide context does not allow efficient initiation on GUG but rather favors initiation on this uAUG or leaky scanning to other initiation codons further downstream, whose resulting translation products can be detected (Figure 5B-C). In fact, all the AUG-like codons tested promote significant leaky scanning. In the presence of the a11wt structure using HEK293FT extracts, the same hierarchy is observed. Comparison between unstressed and stressed HEK293FT cell extracts revealed only minor differences in initiation patterns, although the UUG codon was significantly less efficiently utilized than the AUG (Figure 5D-E). This suggests that stress has a minor effect on the *cis-*regulation of translation initiation on non-AUG codons in our synthetic reporter system.

**Figure 5:**
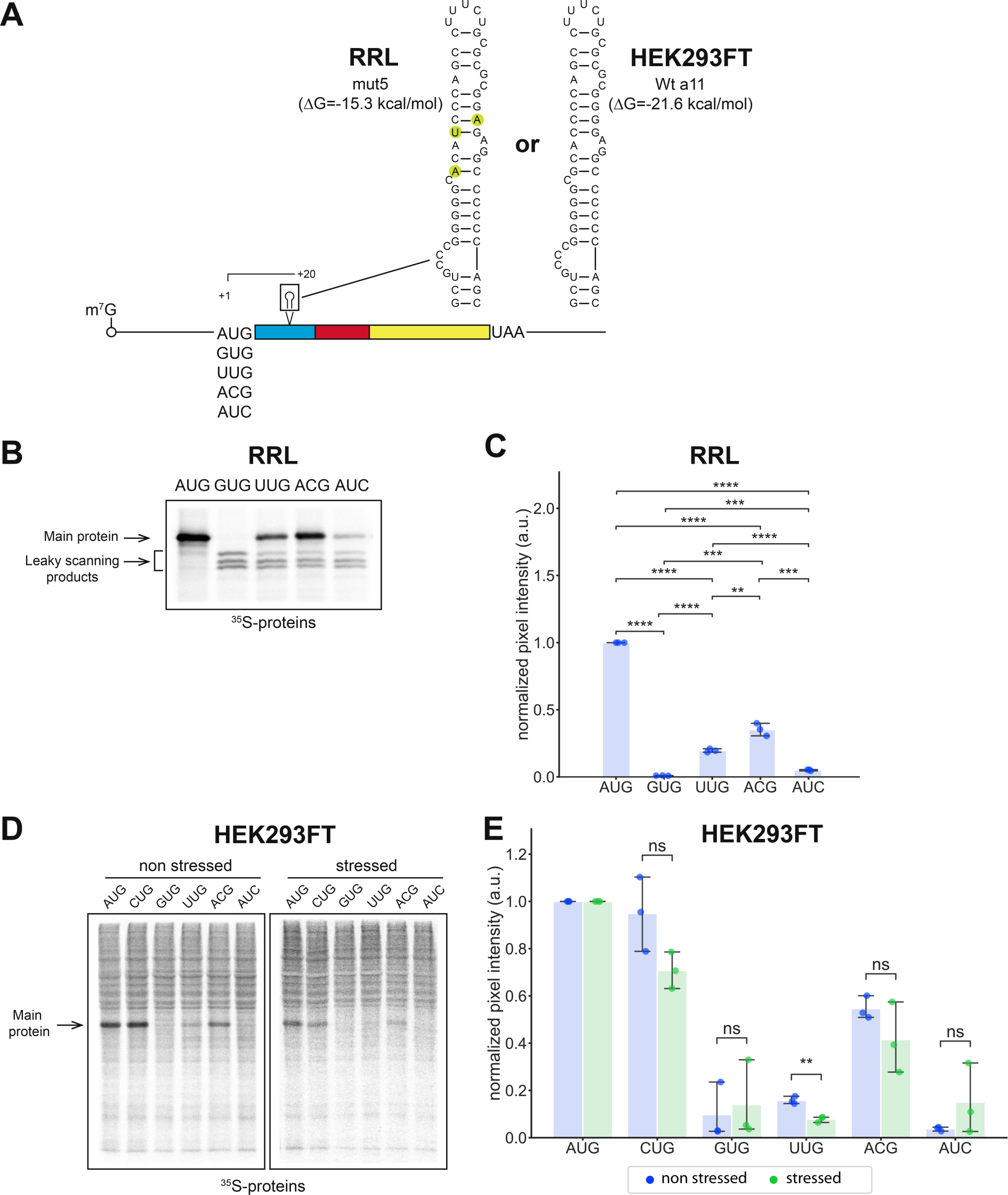
Stable secondary structures do not allow the initiation of translation for all AUG-like codons. **A.** Schematic representation of the RNA reporters and sequences of the stem-loops used during the study. The initiation codons are AUG, GUG, UUG, ACG or AUC codons upstream of the mut5 a11 hairpin for RRL and wt a11 hairpin for HEK293FT which are inserted at +20 position. The nucleotide context of the initiation codons is CGUAAU<u>NNN</u>GAC. **B.** Representative SDS-PAGE experiment of *in vitro* translation products obtained with AUG, GUG, UUG, ACG or AUC codons reporters in RRL. **C.** Quantification of ^35^S incorporation in Renilla luciferase proteins produced in RRL. Bar heights show the mean values with standard deviations indicated. P-values were calculated using a student t-test for independent samples. *: 0.01 < p < 0.05, **: 0.001 < p < 0.01, ***: 0.0001 < p < 0.001, ****: p < 0.0001. **D.** Representative experiment of SDS-PAGE analysis of *in vitro* translation experiment with AUG, GUG, UUG, ACG or AUC codons reporters in unstressed and stressed HEK293FT cell extracts. **E.** Quantification of ^35^S incorporation in Renilla luciferase proteins produced in HEK293FT cell extracts. Results are presented as in C.

Next, we wondered whether the previously established hierarchy of AUG-like codons would be the same using another nucleotide context with the a11wt reporters. For that purpose, the context CGUAAU<u>NNN</u>GAC that was used in previous experiments was changed to GCCACC<u>NNN</u>GCG (Figure 6A). This context is the average nucleotide context of the AUG initiation codon in the human genome, also known as the Kozak sequence ^19,36^. As expected, changing the context rescued initiation on a GUG codon because the Kozak context with a GUG does not contain an out-of-frame AUG codon. However, the GUG codon remains the least efficiently used codon (Figure 6B-C). For the other non-AUG codons, the hierarchy is different from that established with the previous nucleotide context and is consistent whatever the cell-free translation extract used for *in vitro* translation: AUG>UUG≈AUC>>CUG>ACG>GUG.

**Figure 6:**
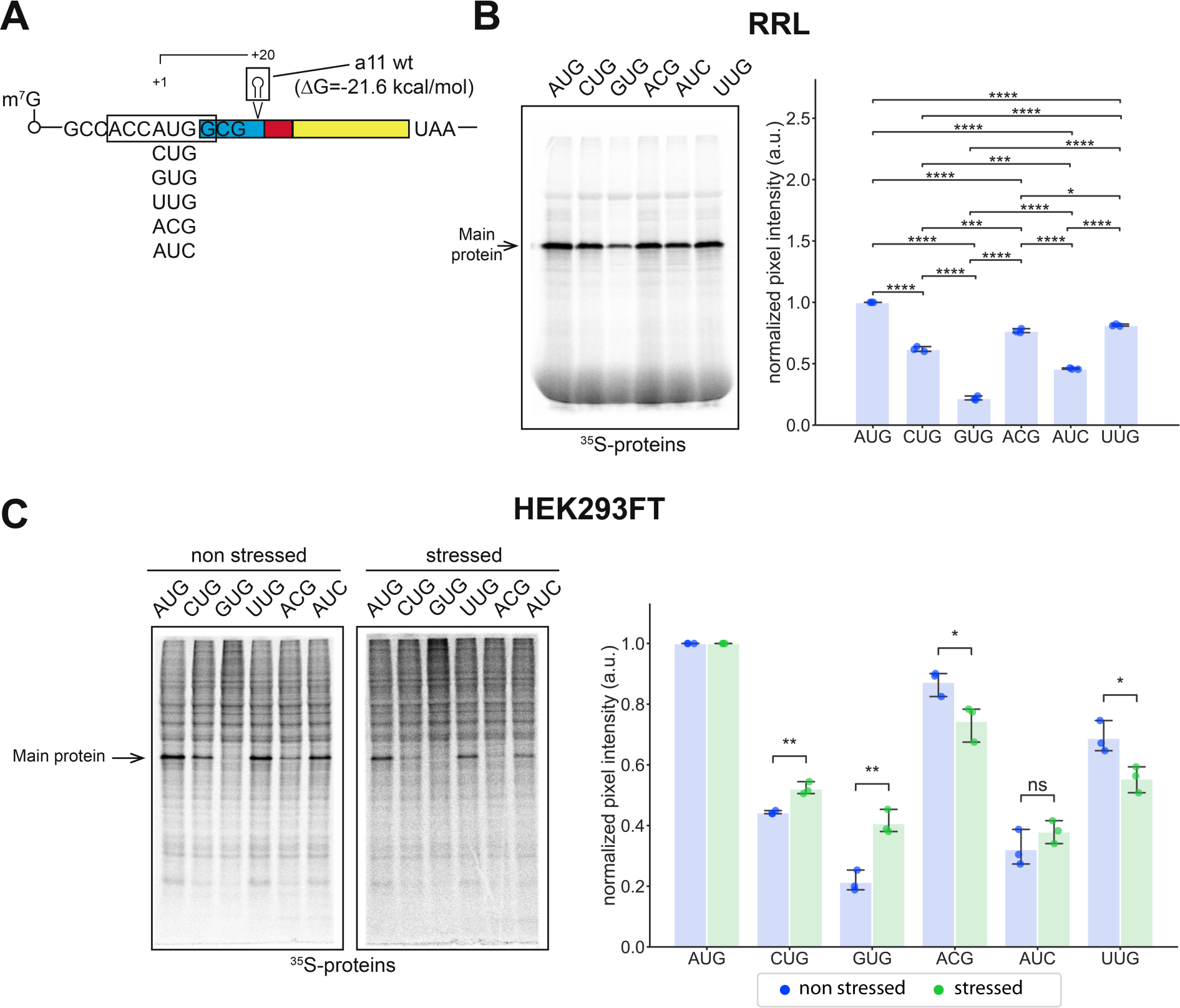
Changing the start-codon nucleotide context influences the efficiency of initiation rescue by START. **A.** Schematic representation of the RNA reporters used in the study. The initiation codons are AUG, CUG, GUG, UUG, ACG or AUC codons upstream of wt a11 hairpin for RRL and HEK293FT which is inserted at +20. The nucleotide context of the tested initiation codons is GCCACC<u>NNN</u>GCG (boxed). **B.** Representative SDS-PAGE of *in vitro* ^35^S radiolabelled translation products obtained with RRL and quantification of pixel intensities of the corresponding gels. Pixel intensities were normalized to those obtained with the AUG reporter. P-values were calculated using a student t-test for independent samples. *: 0.01 < p < 0.05, **: 0.001 < p < 0.01, ***: 0.0001 < p < 0.001, ****: p < 0.0001. **C.** Representative SDS-PAGE of *in vitro* ^35^S radiolabelled translation products obtained with unstressed and stressed HEK293FT extracts and quantification of pixel intensities of the corresponding gels. Pixel intensities were normalized to those obtained with the AUG reporter and are presented as in B. The results obtained with non-stressed extracts are shown in blue and with stressed extracts are shown in green.

### Secondary structures are found downstream of the translation initiation sites of human uORFs

To evaluate the involvement of the START mechanism in non-AUG translation in cells, we investigated genome-wide the presence of RNA structures located at the appropriate distance from start codons. To do that, we used the previously described secondary structure prediction algorithm ^25^ to search for such structures in a set of human small ORFs (smORFs: upstream ORFs, downstream ORFs and “non-coding” ORFs) identified by ribosome profiling ^17^. Briefly, we searched for secondary structures within the +16/+65 window downstream of these ORFs initiation codons in both AUG and non-AUG small ORFs. Among the 5743 recovered smORFs from the dataset of Chothani *et al*., we found rather stable downstream structures (ΔG between -15.6 to -7.1 kcal/mol) in 50% of the analyzed ORFs (Figure 7A). In addition, 50% of these structures have moderate to high predicted stability (more stable than -10.9 kcal/mol) (Figure 7A). These overall statistics suggest that most translated smORFs contain highly stable RNA structures within the +16/+65 window that may facilitate translation initiation. The same analysis carried out according to the start codon shows no obvious correlation, as all codons have similar distributions with rather equal first, second and third quantiles (Figure 7B). We found 556 smORFs starting with an AUG codon and 506 smORFs starting with an non-AUG codon that contain a downstream structure with a stability of at least -15 kcal (Figure 7B, Supplementary File 2 and 3 respectively). Among the non-AUG smORFs, although structures were found with all the non-AUG codons, most of the structures were found in smORFs starting with a CUG (189) and smORFs with a GUG initiation codon (96) (Figure 7C). Only nine structures were found in the smORFS that contain an AUA initiation codon. Altogether, the proportion of non-AUG smORFs containing downstream structures ranges from 7.9% for AUA to 25.1% for ACG (Figure 7C). It also appeared that the average stability of the secondary structures found with the non-AUG start codons is very close to the one found with the AUG codon (Figure 7C). On the other hand, the AUG initiation codon is the most frequently found, accounting for over half of all cases, with a high distribution of highly stable secondary structures (more stable than -10.9 kcal/mol). The second and third most frequent non-AUG codons found are CUG and GUG, respectively, also with a strong distribution of very stable secondary structures. This suggests that the translation of these smORFs could be promoted by these secondary structures. Furthermore, we performed a GO term analysis (Figure 7D), but no explicit term is significantly enriched for transcripts with secondary structure stabilities within the first three quantiles (Supplementary Figure 3), suggesting that all biological processes may be regulated by a START mechanism. However, the fourth quantile (transcripts with secondary structures more stable than -15.6 kcal/mol) shows significantly enriched GO terms linked to the regulation of transcription by RNA polymerase II, signaling cascades, and protein maturation (Figure 7D).

**Figure 7:**
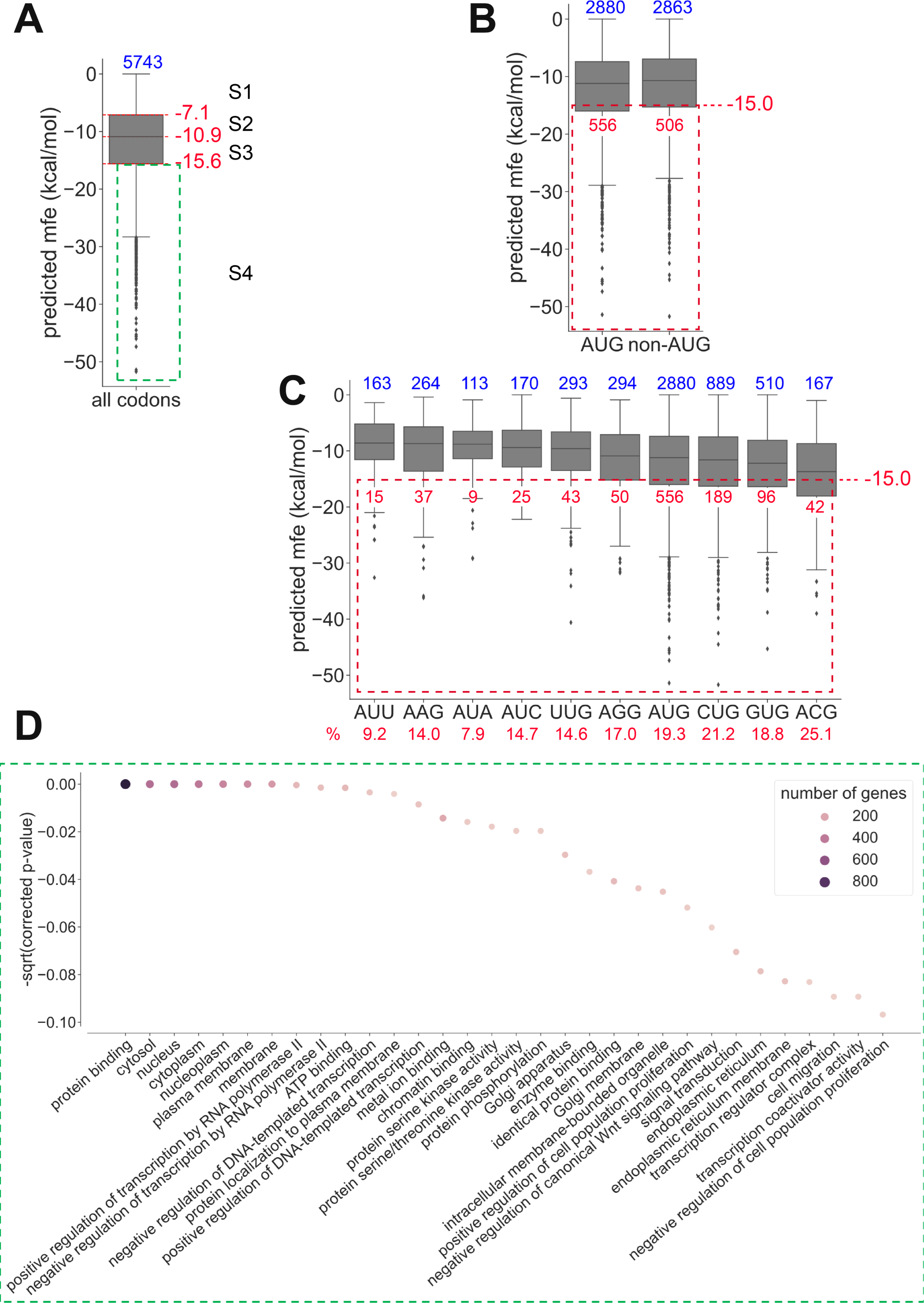
ΔG distributions and GO-term analysis of the predicted secondary structures within the +16/+65 window in the translated human smORFs. **A.** Boxplots showing the **Δ** G (mfe) distributions for all codons, for AUG and non-AUG codons (**B**) and for each codon individually (**C**). Top blue numbers show the population size of each boxplot. Transcripts belonging to each of the four quantiles are labelled S1 to S4, S4 having the transcripts with the most stable structures. The numbers in red show the proportion of smORFs that contain a structure with a stability of at least -15 kcal/mol (shown in a red dashed rectangle in B and C). **D.** GO-terms enrichment analysis of the transcripts belonging to the fourth quantile (S4) of the mfe distribution circled in A by a green dashed rectangle. P-values were calculated with a Fisher exact test, corrected by the Holm method, and scaled with their square root (sqrt).

### Translation of four identified human smORF that initiate at a non-AUG codon

We then arbitrarily chose a few candidates among the transcripts having a smORF starting with a non-AUG codon and a secondary structure more stable than -15 kcal/mol to check whether their downstream structure is indeed required for translation initiation. We chose the -15 kcal/mol threshold stability according to the ones determined by *in vitro* translation assay (Figure 4), although it cannot be excluded that some RNA structures may be differentially modified *in vivo* by specific RNA-binding proteins. The selected transcripts were FAM179B, DDHD2, USP9X and RPTOR. The construction scheme for the corresponding reporter RNAs was similar to the previous ones but with the flanking sequences of the specific smORFs and their own initiation codons (Figure 8A, Supplementary File 1). The non-AUG start codons were AGG, GUG, AUA and CUG. We also designed corresponding mutated structures by introducing destabilizing silent mutations (Figure 8B). The *in vitro* translation patterns were reminiscent of those observed with the mutated versions of the a11 structure. First, we found that the four non-AUG codons tested were efficiently recognized as start codons in the cell-free extracts. Second, translation efficiency in HEK cell extracts is more sensitive to secondary structure stability than in RRL, with drastic decreases in the translation of reporter RNAs with mutated secondary structures (Figure 8C-D). This suggests the involvement of *trans*-acting factors that could be different from one cell extract to another. We also observe that in the case of FAM179B in RRL, the presence of the mutated structure triggers significant leaky scanning, indicating that the wild-type structure is indeed required for the accuracy of translation initiation on AGG codon from FAM179B. In the case of DDHD2, translation in RRL is more efficient when the structure is mutated suggesting that the structure has a negative effect (Figure 8E). Overall, in HEK extracts, initiation on non-AUG codons strictly requires the presence of the wild-type secondary structure (Figure 8F). In addition, there was a marked decrease in the translation efficiency of FAM179B and USP9X RNAs, compared to the efficiencies measured in RRL. Altogether, these experiments suggest that downstream structures do impact the initiation on the AUG-like codons and this effect is modulated by the cell’s content of *trans-*acting factors.

**Figure 8:**
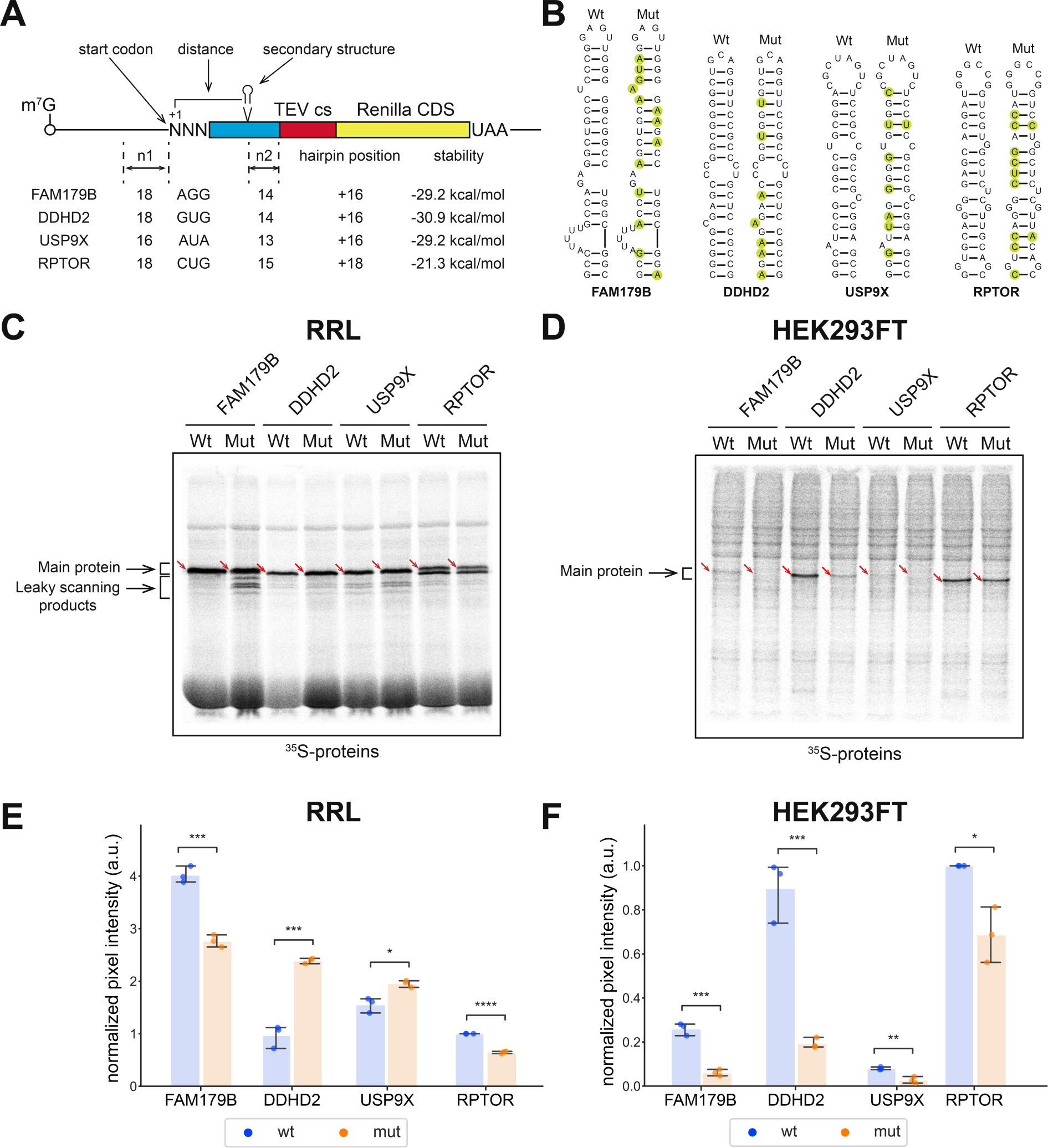
*In vitro* translation of RNA reporters carrying secondary structures from four identified human smORFs using a non-AUG codon. **A.** Schematic representation of the four RNA reporters FAM179B, DDHD2, USP9X, and RPTOR. The stability of the respective structures is indicated as well as the number of nucleotides upstream of the start codon that originates from the human transcript (n1), the start codon, the position of the secondary structure, the predicted stability of the secondary structure and the number of nucleotides downstream the structure that originates from the human transcript (n2). **B.** Representations of the secondary structures of each reporter studied. The silent mutations that were introduced to lower the structure stability are shown in green circles in the Mut stem-loops. **C.** Representative SDS-PAGE of *in vitro* ^35^S radiolabelled translation products obtained with FAM179B, DDHD2, USP9X, RPTOR wild-type and Mut reporters in RRL. The positions of the full-length fusion proteins are shown by red arrows. Lower bands correspond to alternative translation initiation sites resulting from a leaky scanning of the main start codon. In the case of the RPTOR reporter, there is an in-frame AUG at +34. **D.** Representative SDS-PAGE of *in vitro* ^35^S radiolabelled translation products obtained with the same reporters as in C but in HEK293FT extracts. **E-F.** Quantification of ^35^S incorporation in Renilla luciferase proteins produced in RRL and HEK293FT extracts. Bar heights show the mean values with standard deviations indicated. P-values were calculated using a student t-test for independent samples. *: 0.01 < p < 0.05, **: 0.001 < p < 0.01, ***: 0.0001 < p < 0.001, ****: p < 0.0001.

## DISCUSSION

Our data demonstrates that the START mechanism requires an RNA structure with a cell-type dependent minimal stability to promote translation initiation on AUG-like codons and further supports the initial work on that topic ^20^. Considering the length of ribosome footprints provided by toe-printing and ribosome profiling data (approx. 30 nucleotides around the AUG start codon), we expected the optimal distance between the structure and the start codon to be at least 15-16 nucleotides or slightly higher. Indeed, a structure that is too close to the start codon would interfere with the progression of the scanning complex before reaching the start codon. In contrast, a structure located too far away from the start codon would not interact with the scanning complex and consequently lead to leaky-scanning. We found that the optimal distance to favor initiation ranges from 20 to 23 nucleotides downstream of a CUG codon, which is longer than the distance of 14 nucleotides previously described ^20^. As a possible explanation, it can be assumed that the optimal distance also depends on the tri-dimensional shape of the structure. A small stable structure with few base pairs might require a closer distance to the start codon than one with a more sophisticated fold that could for instance create a kink in the structure pointing toward the scanning complex. This possibility requires more investigation. Finally, the sequences of the structure loop itself may also have their own importance as they may interact specifically with either ribosomal components (ribosomal proteins or rRNA) and/or *trans-*acting factors. As an example, such interactions between helix h16 of the rRNA and nucleotides at positions +17 to +19 located immediately upstream of a stable structure have been observed in the case of histone H4 mRNA ^37^.

The START model implies that the stability of the structure (whether or not associated with an RNA-binding protein) must be high enough to stall or at least slow down a scanning complex in order to promote 80S assembly on the codon in the P-site. Therefore we chose as a model RNA structure the a11 GC-rich structure (-21 kcal/mol), as it was previously shown to promote stalling of the scanning pre-initiation complex ^32^. Using this structure as a starting point, we determined that the threshold stability was close to -15 kcal/mol for initiation on CUG codons with RRL, and approximately -20 kcal/mol with extracts from HEK293FT and SHSY5Y cells. These thresholds also allowed initiation on GUG, UUG, ACG and AUC codons, but with distinct efficiencies. Therefore, other parameters are critical such as the nucleotide context, specific *trans-*acting factors, and base-pairing ability with the initiator tRNA^Met^. For instance, changing the start codon nucleotide context resulted in two distinct hierarchies of translation initiation efficiencies among AUG-like codons. Accordingly, previous reports also suggested that initiation on AUG-like codons is influenced by their nucleotide context similarly to AUG codon, with the average preferred nucleotide context being the Kozak context ^36,38,39^.

The AUG-like codon hierarchies we have established using two different start codon nucleotide contexts are rather consistent with the one obtained by ribosome profiling ^15–17^. One striking exception is the GUG codon that is efficiently used in those experiments but not in ours. Thermodynamic studies of the codon-anticodon interaction with a GUG codon demonstrated an increased so-called ‘free-energy penalty’ due to eIF1 and eIF1A interactions that further impairs the initiation on GUG codons but not on CUG codons ^40^. Therefore, initiation on GUG codons is more stringent than on a CUG codon and requires more free energy compensation. Such compensation might come either from a stable-enough downstream secondary structure, an optimal nucleotide environment or from various *trans*-acting factors that modulate the structure stability or start codon selection stringency.

Another example is the balance between eIF5-mimic protein 5MP1/5MP2 and eIF5 that have been shown to respectively increase or decrease the stringency of start codon recognition ^13,41^. Alternatively, Leu-tRNA_CAG_ and Val-tRNA_CAC_ have been described as alternative initiator tRNAs and could be used to initiate translation from CUG and GUG codons respectively under certain stress circumstances through mechanisms that involve non-canonical initiation factors eIF2A or eIF2D ^42^.

Translation initiation on AUG-like codons was assessed in three types of cell extracts, which resulted in distinct required minimal structure stability. First, this suggests that a stable downstream secondary structure is a critical parameter influencing translation initiation on an AUG-like codon. Since the optimal structure stability for efficient initiation on AUG-like codons is dependent on the type of extract used, we further suggest that the involvement of scanning *trans-*acting factors cannot be overlooked. Among these are initiation factors eIF1 and eIF1A which control the codon-anticodon interactions ^5,7^, RNA helicases whose activities were shown to decrease uORFs translation in yeast ^21^ or RNA binding proteins ^22,23^. In particular, the RNA helicase content of the cell would determine the stability range of secondary structures that can be unwound and therefore modulate the use of the START mechanism. A precise assessment of the RNA helicase content in distinct cell types would then be needed to better understand the propensity of these cells to initiate translation from AUG-like codons by a START mechanism.

Along with the determination of the RNA helicase content, secondary structures predictions in 5’UTRs of mRNAs may provide an important tool to predict the use of uORFs with AUG-like codons and unveil potential new ORFs. In a previous study, we demonstrated the broad presence of downstream RNA structures in annotated main open reading frames of various organisms ^25^. Here, we used the same algorithm on the small ORF dataset of Chothani *et al.* 2022 obtained on human primary cells and tissues to analyze downstream secondary structures at alternative translation initiation sites. Among the selected structures, 25% are more stable than -15.6 kcal/mol and half of them are between -7.1 and -15.6 kcal/mol. Moreover, downstream structure stabilities are not correlated with the nature of the start codon. Given that 75% of the identified downstream structures have a moderate to high stability (at least more stable than -7.1 kcal/mol), we hypothesize that the absence of “*cis-* correlation” between the nature of the start codon and downstream structures stability enables fine-tuning of START by *trans-*acting factors depending on the physiological conditions. Furthermore, GO terms enrichment analysis did not show any clear link between transcripts featuring stable downstream structures and subsets of biological processes, although we observed a significant enrichment of GO terms linked to transcription regulation and signaling cascades among transcripts featuring the most stable structures. Overall, this suggests that downstream structures may regulate a broad range of cellular functions.

However, assigning a precise molecular role to these downstream RNA structures requires further investigation. It could be assumed on one hand that some potential start codons have a rather suboptimal nucleotide context that would promote leaky scanning and therefore require a downstream structure to promote initiation. These downstream structures would be modulated by other *trans*-factors that control the structure stability and therefore translation initiation, as it has been observed in yeast ^21^. Integrating these other factors into uORFs translation predictions remains a real challenge, as RNA remodeling is a dynamic process that relies on many factors other than the RNA sequence itself. On the other hand, some of these structures may not be linked to a START mechanism at all, and may play another role for instance in modulating mRNA half-life. Therefore, the percentage of the identified downstream structures that are linked to START remains an open question.

Altogether, we propose that the ubiquity of moderately stable downstream structures at alternative translation initiation sites in human RNAs has evolved as a mechanism for controlling translation initiation through the conditional expression of RNA helicases or RNA-binding proteins, depending on cell-type.

## CONCLUSION

Here, we provide further insights into *cis-*regulatory parameters that influence start codon recognition. Downstream secondary structures can efficiently promote AUG-like translation initiation if they have the required stability to stall a scanning 43S particle. These downstream structures must be located at an optimal distance from this AUG-like codon to trigger codon-anticodon base-pairing in the P site. This is achieved by providing an energetic contribution to the 43S particle with suboptimal AUG-like codons in the P-site. The START mechanism may also rescue translation initiation on start codons with a suboptimal nucleotide context that would otherwise lead to leaky scanning. A better understanding of how frequently this atypical initiation mechanism is used in living cells might help to predict more accurately translation initiation sites in cellular mRNAs and give a more precise picture of the complete proteome. Overall, adjusting the stability of RNA structures downstream of potential start codons by the expression level of specific RNA helicases or RNA-binding proteins in various cell types is an elegant way to turn on and off translation initiation on specific codons depending on the physiological conditions.

## ACKNOWLEDGMENTS

This work is funded by *Agence Nationale pour la Recherche* (ANR-17-CE12-0025-01, ANR-17-CE11-0024, ANR-20-COVI-0078, ANR-21-CE12-0028-01, ANR-21PRRD-0001-01), by *Fondation pour la Recherche Médicale* (project CoronaIRES and ALZ201912009641), by *Fondation Bettencourt,* by University of Strasbourg and by the *Centre National de la Recherche Scientifique*. We would also like to thank Dr. Sebastian Schafer for kindly providing us with DNA sequences of the 7,767 high-confidence smORFs detected in the human genome ^17^.

## AUTHOR CONTRIBUTIONS

AT designed the experimental strategy of this study, performed most of the experiments, interpreted the data and wrote the manuscript. FA performed the previous experiments on hox a11 that enabled this study and proofread the manuscript. LD created and used the scripts required for the screening of the smORF database, GE interpreted the data and participated to the writing of the manuscript, FM designed the experimental strategy, supervised the conducted experiments, interpreted the data and wrote the manuscript.

## DATA AVAILABILITY

The authors confirm that the data supporting the findings of this study are available within the article and */* or its supplementary data.

