## Supplementary material for "Critical *cis*-parameters influence STructure Assisted RNA Translation (START) initiation on non-AUG codons in eukaryotes"

#### SUPPLEMENTARY INFORMATION

**Supplementary Table 1: oligonucleotides' sequences**

**Supplementary File 1: sequences of the reporter RNAs used in this study**

**Supplementary File 2: downstream RNA structures data in the recovered human smORFs sequences starting with an AUG codon**

**Supplementary File 3: downstream RNA structures data in the recovered human smORFs sequences starting with a non-AUG codon**

**Supplementary File 4: The Python module named toolbox.py is imported by some of these scripts (Supplementary File: Python_scripts.zip).**

**Supplementary Figure 1: the cell-free translation extracts used in this work are cap-dependent.**

$m^7G$- or A-capped 5'UTR β-globin Renilla reporters (**A**) were *in vitro* translated in RRL or in HEK cell lysates. Renilla quantification shows that our extracts are mostly cap-dependent, with A-capped translation being 30% of $m^7G$ translation in RRL (**B**) and 20% in HEK cell lysates (**C**, in blue). Interestingly, the HEK extracts prepared from stressed cells (**C** and **D**, in green) are twice less cap-dependent, with A-capped translation being 40% of $m^7G$ translation.

**Supplementary Figure 2: functional characterization of the HEK cell-free translation extracts.**

We prepared cell-free translation extracts from HEK293FT cells that were grown in optimal conditions and in stress conditions. We checked that the stress induced by DTT exposure induced eIF2$\alpha$ phosphorylation by western blot (**B**). The phosphorylation rate of eIF2$\alpha$ is two times higher in stressed cells compared to unstressed cells. We tested these extracts for *in vitro* translation with two reporters (**A**): a luciferase uncapped RNA reporter containing in its 5'UTR the IRES of the intergenic region (IGR) from the Cricket Paralysis Virus (CrPV) to test cap-independent translation and a canonical cap-dependent reporters. As expected, cap-dependent translation is strongly inhibited in the stressed extracts while IGR-driven translation is significantly enhanced. IGR does not require translation initiation factors and is therefore not sensitive to eIF2$\alpha$ phosphorylation. Since canonical translation is aborted, the pool of available ribosomes in these extracts is higher which most probably explains the increase of translation by IGR. Altogether, we concluded that these extracts are suitable for the characterization of *cis*-regulatory elements involved in both AUG and non-AUG translation.

**A.** $m^7G$ and IGR translation efficiencies rely on the stress level of the HEK extracts

$^{35}S$ signal quantification of *in vitro* translated Renilla luciferase with HEK293FT (blue) or stressed HEK293FT (green) cell-free translation extracts from $m^7G$- or IGR- driven translation of the reporter RNAs shown on the left. Error bars represent the 99% confidence interval calculated from three independent replicates using the t-distribution. p-values were calculated using a student t-test for independent samples. cds: coding sequence. *: $0.01 < p < 0.05$, **: $0.001 < p < 0.01$, ***: $0.0001 < p < 0.001$, ****: $p < 0.0001$

**B.** Calculation of the stress level of HEK293FT cell-free translation extracts by Western Blot measurements of eIF2 phosphorylation. To determine the initial rate (at t = 0 min) of phosphorylated eIF2α (eIF2α-P) in both extracts, we hypothesized that the phosphorylation level of eIF2α is 100% after 75 min of *in vitro* translation. Consequently, the level of the eIF2α-P is considered equal to total eIF2α after 75 min. Then, the initial rate of eIF2α-P is calculated using the ratio eIF2α-P (t = 0 min) / eIF2α-P (t = 75 min).

Left: Western Blot images with antibodies specific for non-phosphorylated eIF2α (bottom) or phosphorylated eIF2α (top) using cell-free translation extracts prepared from HEK293FT cells cultured in physiological conditions (HEK-R, blue) or in stressed conditions (HEK-S, green)

Right: quantification of eIF2α phosphorylation using the Western Blot images

The dashed lines indicate the images (each in one black rectangle) were cropped (*i.e.* the lanes are not adjacent).

**Supplementary Figure 3: ΔG distributions and GO-term analysis of the predicted secondary structures within the +16/+65 window in the translated human smORFs.**

The boxplot shows the ΔG (mfe) distributions for all codons (**A**). Transcripts belonging to each of the four quantiles are labelled S1 to S4, S4 having the transcripts with the most stable structures. Top blue number shows the population size of the boxplot. (**B-E)** GO-terms enrichment analysis of the transcripts belonging to the first (**B**), second (**C**), third (**D**) and fourth quantiles (**E**) of the mfe distribution (**A**). P-values were calculated with a Fisher exact test, corrected by the Holm method, and scaled with their square root (sqrt).

#### Supplementary Figure 1

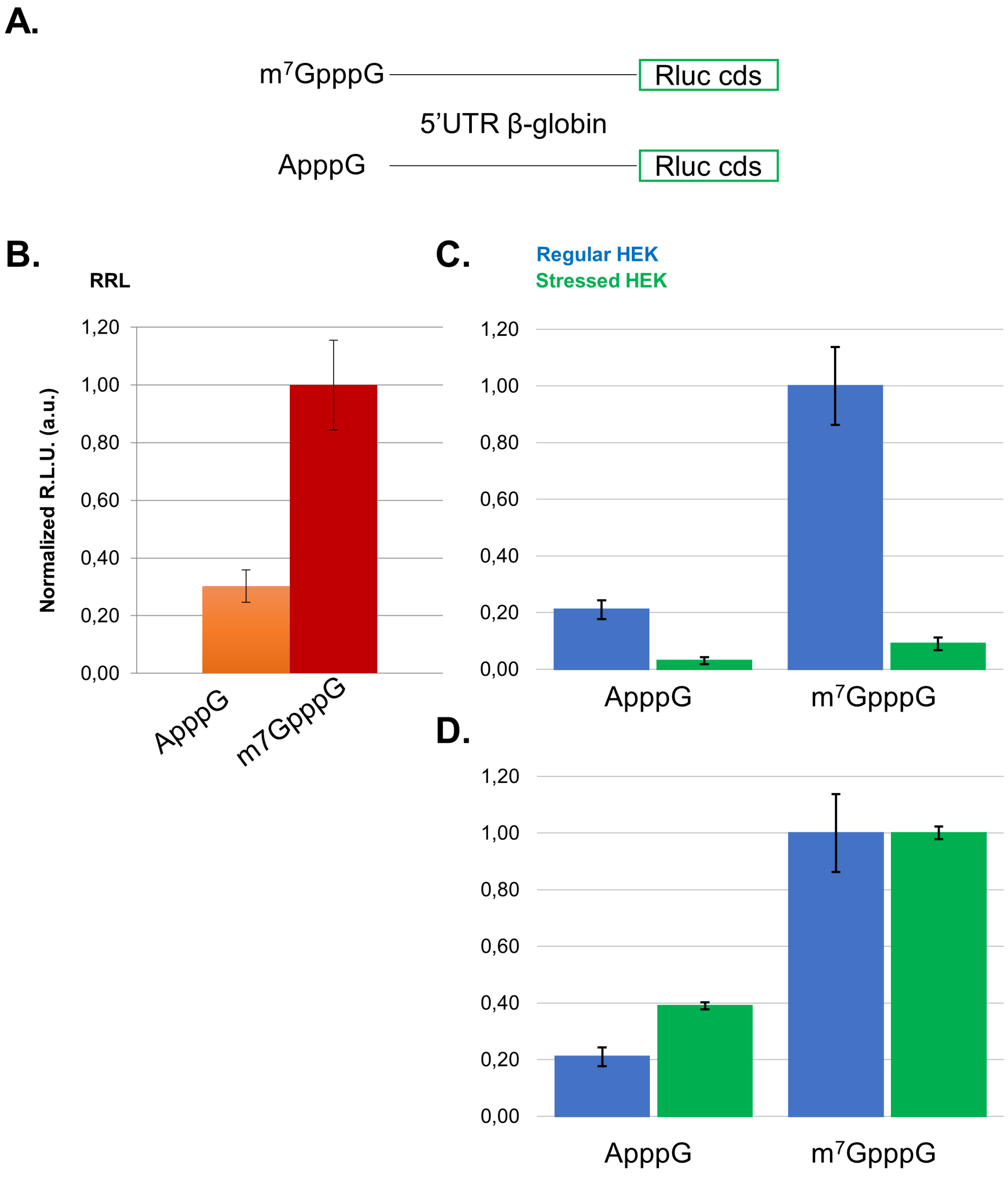

#### Supplementary Figure 2

**A.**

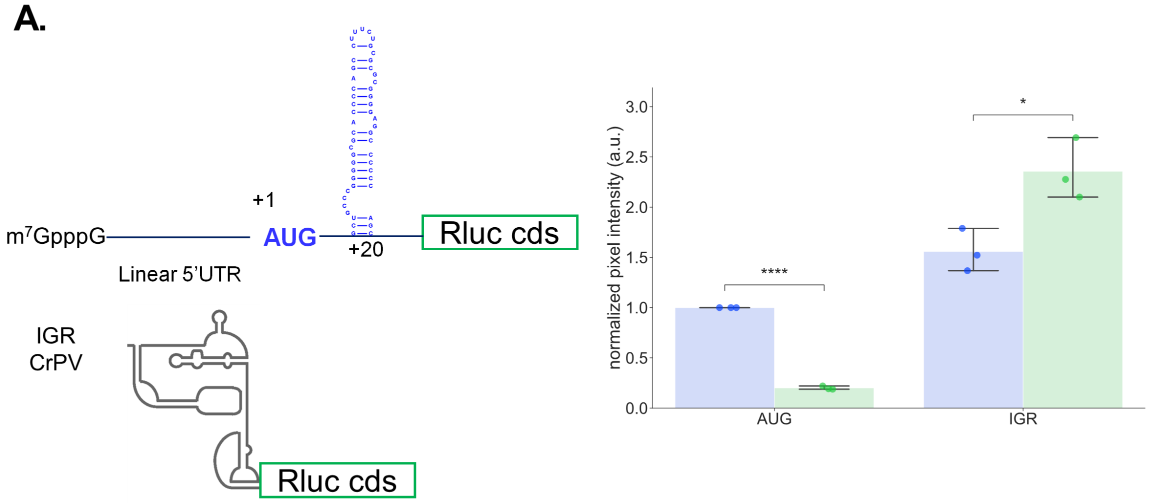


**B.**

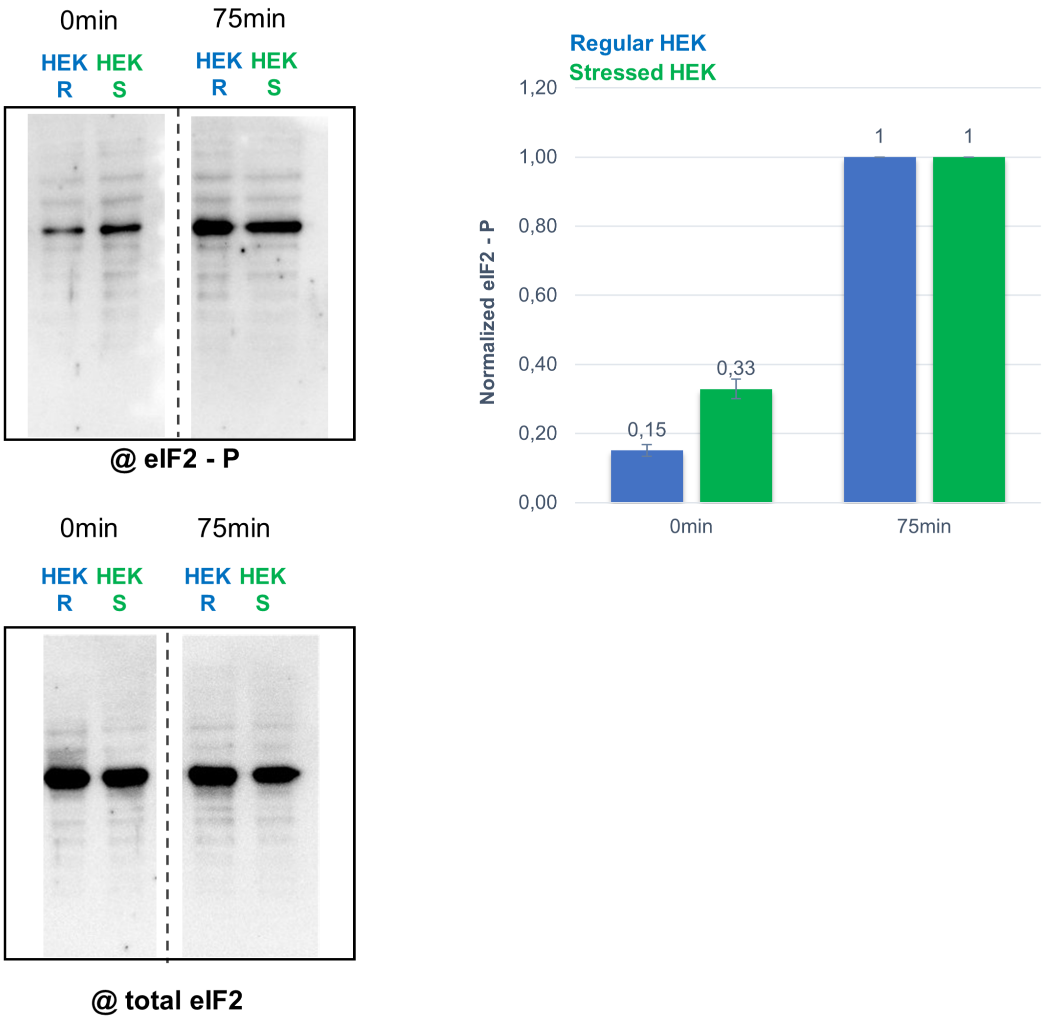

#### Supplementary Figure 3

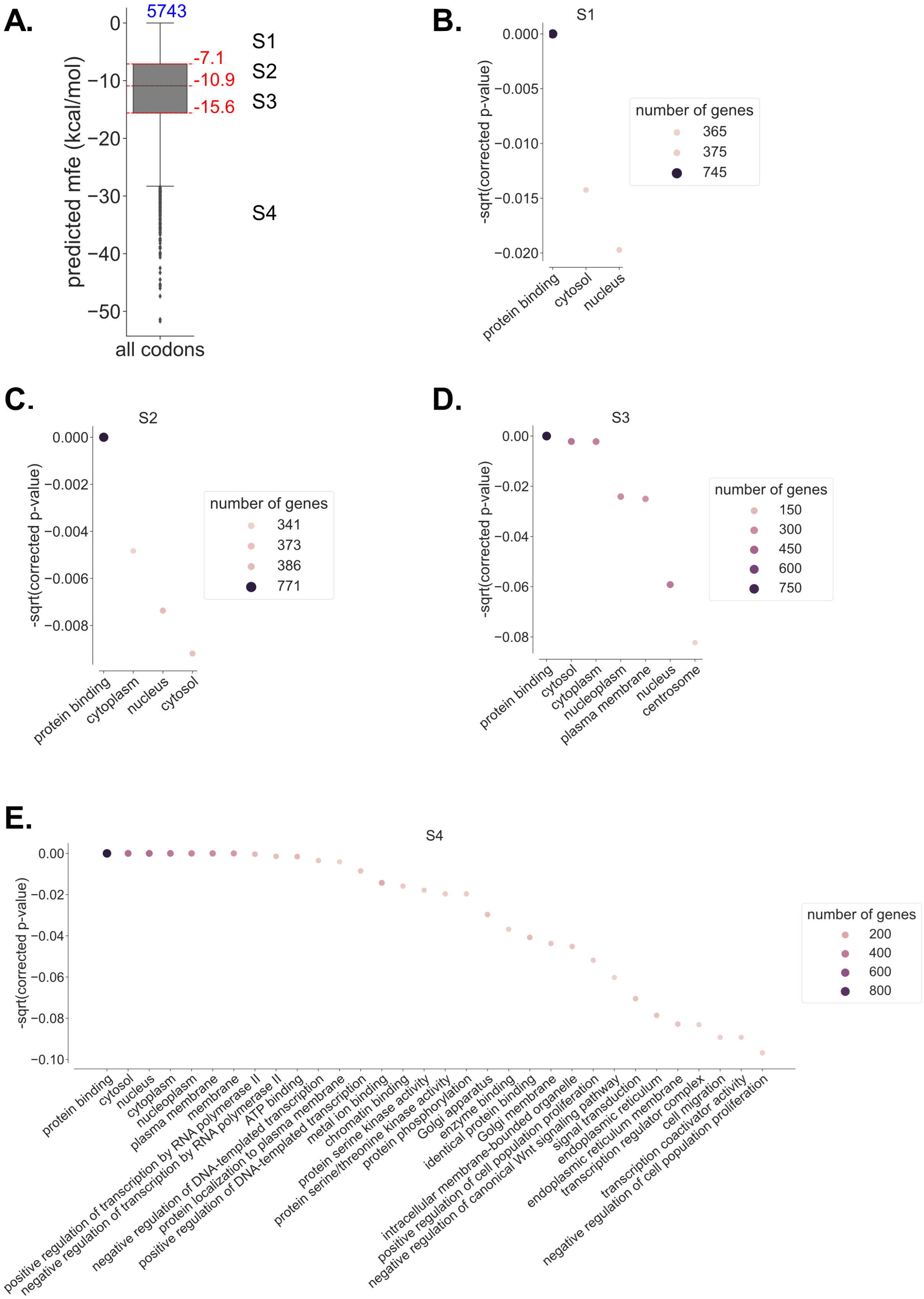

### Supplementary Table 1

| n° in text | Primer name | Sequence 5'-3' |
|---|---|---|
| 1 | fw_T7-RNA-158 | CAACAAATATTAATACGACTCACTATAGGGAAAAACACGACCATAATC |
| 2 | fw_T7-IGR | CAACAAATATTAATACGACTCACTATAGGCAAAAATGTGATCTTGCTT |
| 3 | rev_Renilla_pUC19 | GCATGCCTGCAGGTCGACTACTAGTTTATTGTTCATTTTTGAGAAC |
